# Diversity of culturable endophytic fungi associated with avocado orchards under organic and conventional management

**DOI:** 10.1101/2025.01.31.635834

**Authors:** Daniel Sánchez-Hernández, John Larsen, Frédérique Reverchon

## Abstract

Endophytic fungi constitute a great resource for the implementation of sustainable agricultural practices. However, their diversity may be altered by agrochemicals used in conventionally managed agrosystems. Here we explored the culturable diversity of endophytic fungi associated with avocado (*Persea americana* Mill.) in conventional and organic orchards from three counties of the world’s leading productive region. In total, 529 endophytic fungal isolates were obtained from branches and roots of avocado trees. Identification based on ITS sequencing showed that most isolates belonged to the Ascomycota phylum (98 %) and were grouped into 52 fungal genera, representing 102 tentative species. The endophytic fungal community was dominated by genera which were common to both management types, namely *Fusarium, Colletotrichum, Dactylonectria*, and *Diaporthe* / *Phomopsis*. Twenty fungal genera were exclusive to organic orchards (e.g., *Hypoxylon* and *Talaromyces*) and 10 were only found in conventional orchards (e.g., *Trichoderma*). Fungal species richness and diversity were higher in organic than in conventional orchards. Ordination and clustering analyses revealed a stronger effect of site than of management type on endophytic fungal community structure, highlighting the influence of local factors in shaping the culturable endophytic fungal community. Overall, our findings confirm the negative impact of conventional management on the diversity of crop-associated endophytic fungi and the significant influence of local factors on endophytic fungal assemblages. Moreover, our results emphasize the need to assess the risk that possible latent fungal pathogens may represent for the crop or dominant forest species surrounding avocado orchards.

## Introduction

Endophytic fungi are a widespread group of fungi that colonizes the internal tissues of plants without causing disease symptoms [1]. Previous studies have suggested that they are recruited from the soil, penetrating plants through natural openings following complex molecular signaling processes [2]. The species composition of endophytic fungal communities changes depending on the host plant species, tissue type, age, and abiotic factors such as climatic conditions [3–5]. Endophytic fungi exhibit various types of interactions with their hosts, ranging from latent pathogens, commensals, mutualists, temporary residents, to saprobes [6, 7]. Some endophytic fungi have been documented to induce biochemical and physiological changes in plants, enhancing their tolerance to salinity stress, heavy metals, and xenobiotics such as different pesticides [8–11]. In addition, they can promote plant growth by improving their nutrient acquisition, inducing indole production [12] and protecting them against pests and diseases [13–15].

Given their significant role for plant health and productivity, endophytic fungi constitute a great microbial resource for sustainable agricultural management. However, their diversity may be threatened by the indiscriminate use of agrochemicals or other conventional agricultural practices [16–18]. The influence of agronomic management on the structure and diversity of endophytic fungi has been investigated in several crops. da Costa-Stuart *et al*. [19] evaluated the effects of organic and conventional management on the community of endophytic fungi associated with soybean (*Glicine max* (L.) Merr.). Their results show that the diversity, richness and evenness of endophytic fungal communities decreased in conventional farming with respect to organic management, and that the applications of pesticides induced a larger dominance of the *Alternaria* genus, a potential pathogen, within the community. These findings were consistent with those from [20], who found a larger abundance and diversity of endophytic fungi associated with organic farming of maize (*Zea mays* L.), tomato (*Solanum lycopersicum* L.), pepper (*Piper nigrum* L.) and watermelon (*Citrullus lanatus* Thunb.), than in conventional cropping systems. In general, these results highlight the negative impact of intensive agricultural management on crop-associated endophytic fungi.

Avocado (*Persea americana* Mill.) is a tree crop belonging to the Lauraceae family, highlighted for its culinary value and significant economic relevance at a global level [21]. Currently, Mexico leads the world production and export of this fruit, reaching a planted area of 218,821 hectares in 2024 and an annual estimated production of 2,652,501 tons [22]. However, the productive success of avocado has led to extensive deforestation and land use change [23], contamination by agrochemicals, and loss of biodiversity and biotic resources, with negative consequences at the environmental, social and human health levels [24–26]. In this context, endophytic fungi may constitute a promising alternative to agrochemicals, to mitigate the negative environmental impact of avocado production whilst sustaining its productivity.

Few studies have investigated the diversity of endophytic fungi associated with avocado trees, focusing mainly on isolating *Trichoderma* strains. In South Africa, Hakizimana *et al*. [27] isolated 24 species of endophytic fungi from avocado roots and showed that endophytic fungi of the genus *Trichoderma* exhibited antagonistic activity against the pathogen *Phytophthora cinnamomi*. In Florida, Shetty *et al*. [28] revealed that fungal endophytes of avocado branches differed between organic and conventional production systems, suggesting a significant influence of pesticides, and highlighted a large proportion of potential pathogenic taxa within the endophytic community. In Mexico, Andrade-Hoyos *et al*. [29] isolated and validated the biocontrol activity of ten endophytic strains of *Trichoderma* associated with avocado roots against *P. cinnamomi*, observing a 50 % decrease in the incidence of the disease in plants inoculated with *T. harzianum* and *T. asperellum*. The biocontrol potential of endophytic *Trichoderma* strains was also reported by López-López *et al*. [30], who highlighted the antifungal activity of strain *T. harzianum* TSMICH7 against the plant pathogens *Neofusicoccum parvum*, *Colletotrichum gloeosporioides*, *Diaporthe* sp., and *Phomopsis perseae* in avocado fruits, suggesting its potential use as a biological control agent in postharvest disease problems. More recently, Nieves-Campos *et al*. [12] isolated 89 endophytic fungi from the roots of avocado nursery plants, which were grouped into 24 morphotypes of five genera: *Fusarium*, *Penicillium*, *Metapochonia*, *Mortierella*, and *Alternaria*. Subsequent screenings revealed that isolates from 20 of the 24 morphotypes could successfully inhibit the growth of *P*. *cinnamomi in vitro*. Moreover, strains from the *Metapochonia* and *Mortierella* genera were identified as growth promoters of *Arabidopsis thaliana* in *in vitro* co-inoculation assays. These findings show that avocado endophytes may harbor great potential as biocontrol agents or biofertilizers, although their diversity has not yet been harnessed. Furthermore, how agricultural practices could impact the diversity of avocado endophytes is still unknown. Our objective was thus to explore the culturable diversity of endophytic fungi associated with avocado roots and branches in conventional and organic orchards of the world’s top productive region.

## Materials and methods

### Sampling and isolation of endophytic fungi

During April and May 2023, root and branches samples were collected from one organic and one conventionally managed avocado orchard (cv. Hass) in the counties of Uruapan, Tancítaro and Tacámbaro, located in the state of Michoacán, Mexico (**Supplementary Material S1**). These counties were selected because they were the ones with the largest cultivation area in the state in 2020 [31]. Specific details of the sampling sites are presented in **Supplementary Material S2**. All organic orchards were certified and adhered to restrictions that prohibit the use of synthetic fertilizers and pesticides not authorized in organic agriculture. On the other hand, orchards under conventional management used permitted chemical products, including fertilizers and pesticides listed in the basic table of products authorized for export, in accordance with current regulations (personal communication with farmers).

Tissue sampling from avocado trees was carried out by selecting six asymptomatic trees per orchard. From each tree, four samples of branch tissue of approximately 1-2 cm in diameter were collected, as well as four samples of secondary roots. All samples (branches and roots) were taken in different zones of the tree according to the four cardinal points. The collected samples were placed in plastic bags, labeled, transported in coolers and stored at 4 °C for subsequent laboratory processing.

The isolation of fungal endophytes was carried out at a maximum of 48 h after sampling according to the methodology described by Hakizimana *et al*. [27]. Branches and roots were first washed with common detergent in running tap water, then 3-cm segments were cut and immersed in 70 % ethanol for 1 min, in a 2 % sodium hypochlorite solution for 5 min and rinsed five times with sterile distilled water. Once the surface disinfection was carried out, segments were further cut in 0.5-cm pieces and plated in two different culture media: potato dextrose agar (PDA) supplemented with chloramphenicol at 50 mg L^-1^, and oat agar with CTAB (cetyl-trimethyl-ammonium bromide) as a selective medium for entomopathogenic fungi (oat 60 g L^-1^, agar 15 g L^-1^ and CTAB 600 mg L^-1^). For each medium, three branch samples or three root samples from the same tree were plated. As a control, an aliquot of 200 µl of the last rinse was streaked in a Petri dish on the PDA + chloramphenicol culture medium, to verify the success of the surface disinfection process. Subsequently, the plates were incubated at 25 °C for 15 days in dark conditions, and fungal growth was monitored daily. At seven and 14 days, subcultures of the fungi that emerged from the tissues were performed.

### Morphological and molecular identification

The morphology of fungal isolates was examined macroscopically, focusing on colony characteristics such as color, topography, pigmentation on the front and back of the culture medium, texture and size [32, 33]. Based on these characteristics, isolates were grouped into morphotypes for each of the sampled orchards.

One fungal isolate per morphotype was randomly selected for molecular identification. Fungal DNA extraction was carried out following the methodology by Izumitsu *et al*. [34], with slight modifications. The fungi were grown in PDA for 7 days at 25 °C, after which a portion of the mycelium was taken with a sterile dissection needle and suspended in 100 µl of 1X TE buffer. Micro-tubes with the mycelium samples were placed in microwave for 50 s (1000 w) followed by 30 s of incubation at room temperature, subsequently microwaved again for 50 s (1000 W) incubated at -20 °C for 10 min, and centrifuged at 13,000 rpm for 5 min. The supernatant served as a template to perform the PCR.

Partial amplification of the 18S rRNA gene was carried out using the universal primers ITS1F (5’-CTTGGTCATTTAGAGGAAGTAA-3’) and ITS4 (5’- TCCTCCGCTTATTGATATGC-3’). The amplification was performed in a volume of 25 µl, with 13.7 µl of sterile distilled water, 2.5 µl of 5X colorless buffer, 2.5 µl of MgCl_2_ (25mM), 1.0 µl of deoxyribonucleotide triphosphate (dNTPs, 10mM), 1.0 µl of each primer (10 µM), 0.1 µl 1% BSA, 0.2 µl GoTaq® DNA polymerase (Promega), and 3 µl DNA template. The optimized PCR thermal profile was 94 °C for 3 min, followed by 30 cycles, each containing a denaturation step at 94 °C for 30 s, a hybridization step at 50 °C for 30 s, and final extension at 72 °C for 7.0 min. The DNA amplicons were purified using the Wizard® SV Gel Cleaning System and PCR kit (Promega) following the manufacturer’s instructions and subsequently sent for sequencing at Macrogen, Inc., South Korea. The obtained sequences were deposited in GenBank under accession numbers PQ684493-PQ684779 and PQ687555.

### Phylogenetic and statistical analyses

Sequences were manually checked in BioEdit 7.7 [35]. Once the sequences were edited, the search for the closest species was carried out using the BLASTn algorithm with the GenBank nucleotide database (www.ncbi.nlm.nih.gov). Relative abundances were calculated using the *dplyr* package in R [36]. The stacked bar graphs used to visualize these abundances were created with the *ggplot2* package [37] in RStudio [38]. A Venn diagram was constructed using the online tool provided by Bioinformatics and Evolutionary Genomics (www.bioinformatics.psb.ugent.be/webtools/Venn/).

Diversity analysis based on Hill’s numbers was conducted with the *iNEXT* package in R [39] and the results were visualized as boxplots created with *ggplot2* in RStudio. The normality of the diversity estimates was assessed using the Shapiro-Wilk test. The effects of management practices (organic and conventional) and geographic location (counties) on fungal community diversity metrics were evaluated using a two-way Analysis of Variance (ANOVA), followed by a Tukey post-hoc test (*p* ≤ 0.05) in RStudio [38]. Non-Metric Multidimensional Scaling (NMDS) based on the Bray-Curtis distance matrix and the construction of dendrogram were performed in R using the *vegan*, *ggplot2*, *ggforce* and *ggdendro* packages in RStudio [40–42].

## Results

### Isolation and morphotyping of endophytic fungi

A total of 529 endophytic fungal isolates were obtained from the branches and roots of avocado trees across all orchards, with a nearly equal proportion of isolates derived from roots (52.93 %) and branches (47.07 %). Of these, 246 isolates were obtained from conventional orchards, while 283 were retrieved from organically managed orchards (**Fig. 1**). These isolates were grouped into 291 morphotypes by morphological characterization, separating fungi with similar morphology but retrieved from different orchards. Of these 291 morphotypes, 122 corresponded to conventional orchards and 169 to organic orchards.

**Figure 1.**
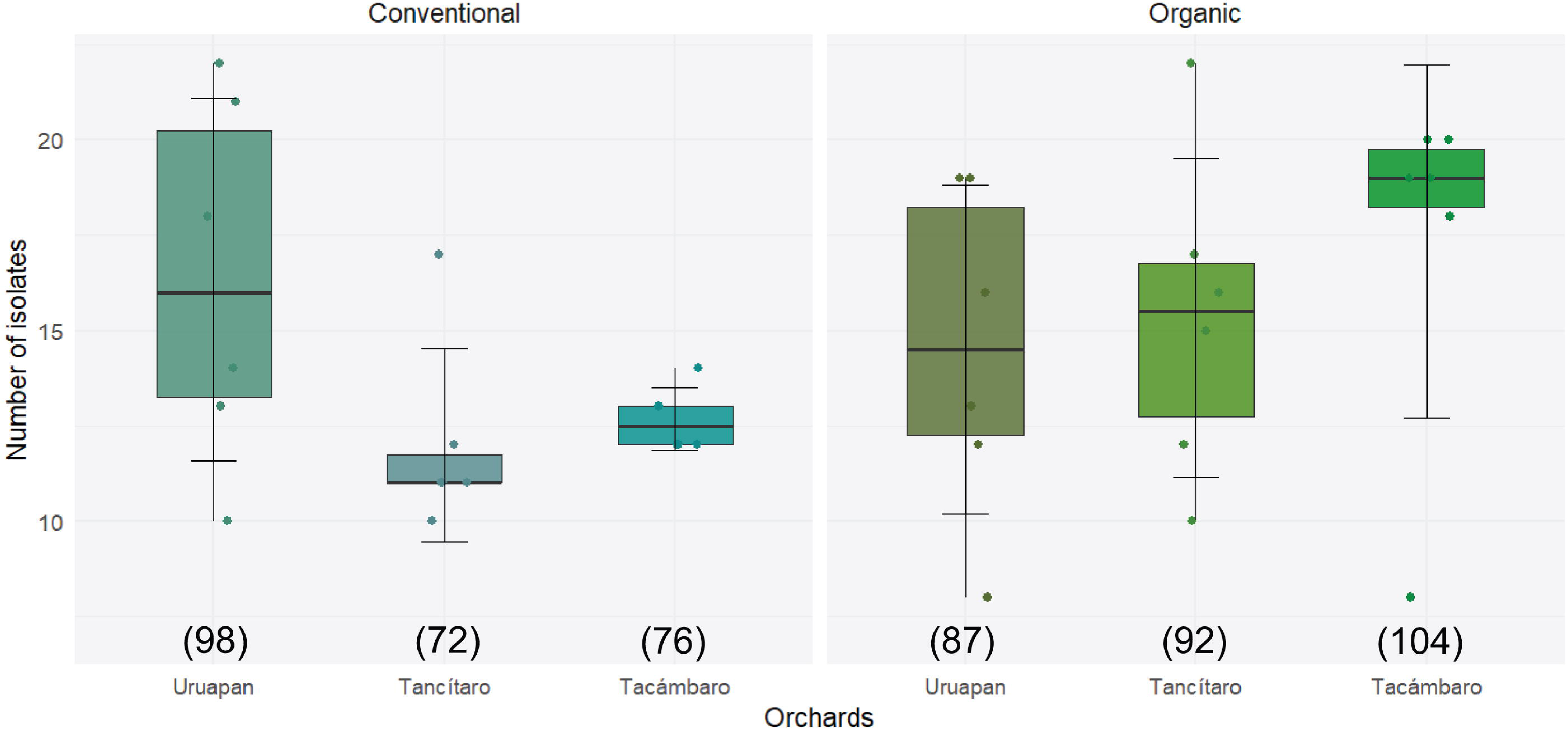
Number of fungal isolates retrieved in each sampled avocado orchard. The left panel shows the number of isolates obtained in three conventionally managed orchards, while the right panel displays the corresponding data for three organically managed orchards. Scatter points represent the number of fungal isolates obtained per tree (*n* = 6). No significant differences were found in the number of isolates between orchards (ANOVA, *p* ≤ 0.05). The total number of isolates recovered per orchard is indicated below each respective box plot.

### Identification of the isolated avocado fungal endophytes and species composition

The molecular identification of one representative isolate per morphotype revealed that these 291 morphotypes represented 52 genera and 102 tentative species (**Supplementary Material S2**). These 52 fungal genera belonged to two phyla: Ascomycota (subphyla Pezizomycotina, 98.1 % of genera) and Basidiomycota (subphyla Agaricomycotina, 1.9 %). These two groups were divided into five classes, the two most dominant classes being Sordariomycetes (61.5 %) and Dothideomycetes (30.8 %). At the order level, the isolates were grouped into 14 defined orders and one *Incertae sedis* (*inc. sed*.), with 29 defined families and three *inc. sed*., groups as detailed in **Fig. 2**.

**Figure 2.**
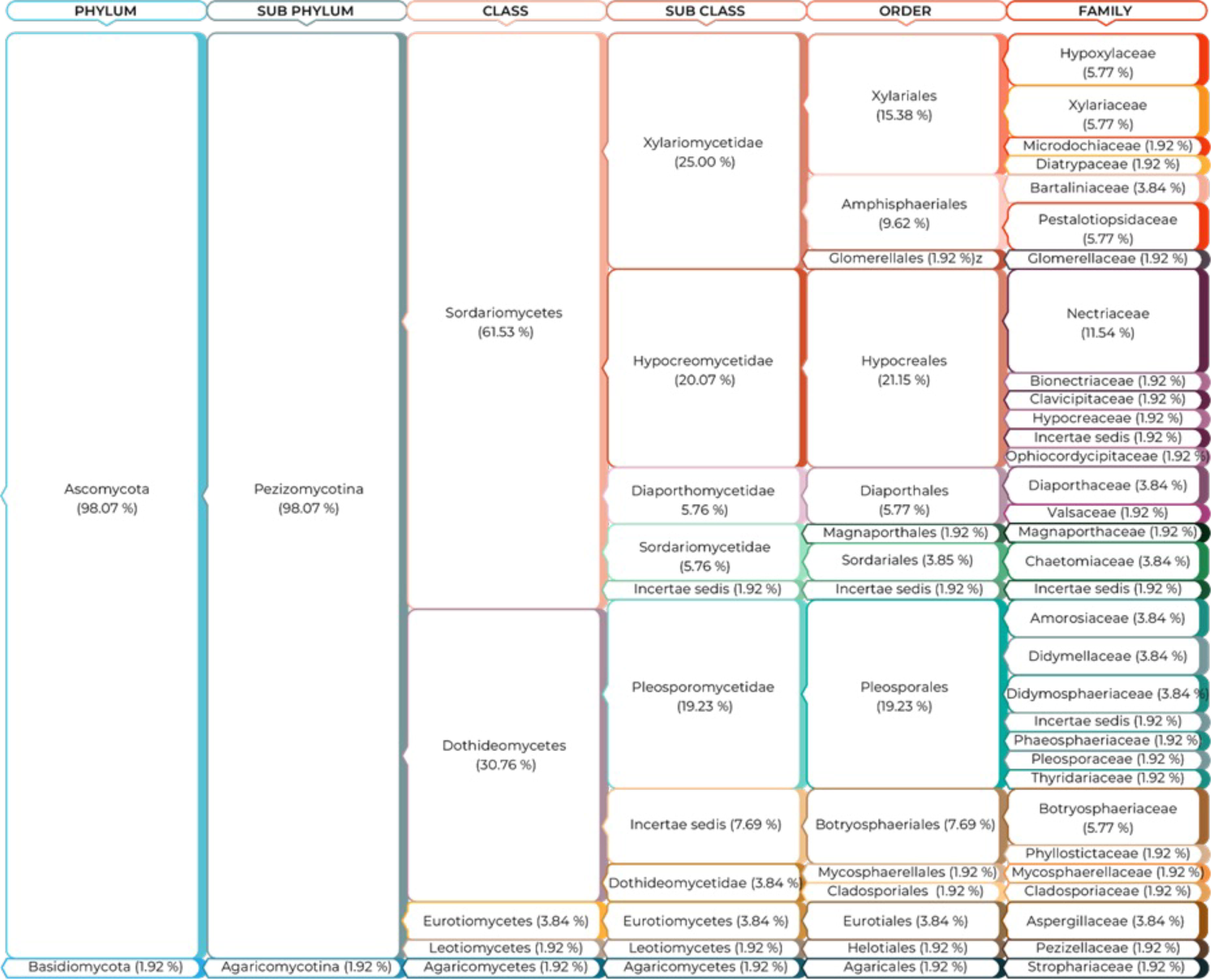
Taxonomic composition of the endophytic fungal community isolated from branches and roots of avocado trees in conventional and organic orchards. The relative distribution of morphotypes is shown at each taxonomic level, from Phylum to Family. Made with information retrieved from the Index Fungorum 2024 (www.indexfungorum.org).

**Figure 3** represents the relative abundances of the identified genera and their distribution across the production systems and counties. Of the total number of isolates, 56.43 % were represented by genera present in all orchards, independently from their management system: *Fusarium* was the most abundant genus, accounting for 23.5 %, followed by *Colletotrichum* (18.6 %) and *Dactylonectria* (14.4%). Ten fungal genera with moderate abundance summed 30.1 % and included *Diaporthe* (6.8 %), *Phomopsis* (anamorph state of *Diaporthe*, 5.7%), *Penicillium* (5.1 %), *Idriella* (3.4 %), *Cladosporium (*2.7 %), *Didymella* (1.5 %), *Neofusicoccum* (1.3 %), *Xylaria* (1.3 %), *Alternaria* (1.1 %) and *Trichoderma* (1.1 %). The remaining 39 genera each accounted for less than 1.0 % of the total (**Fig. 3**).

**Figure 3.**
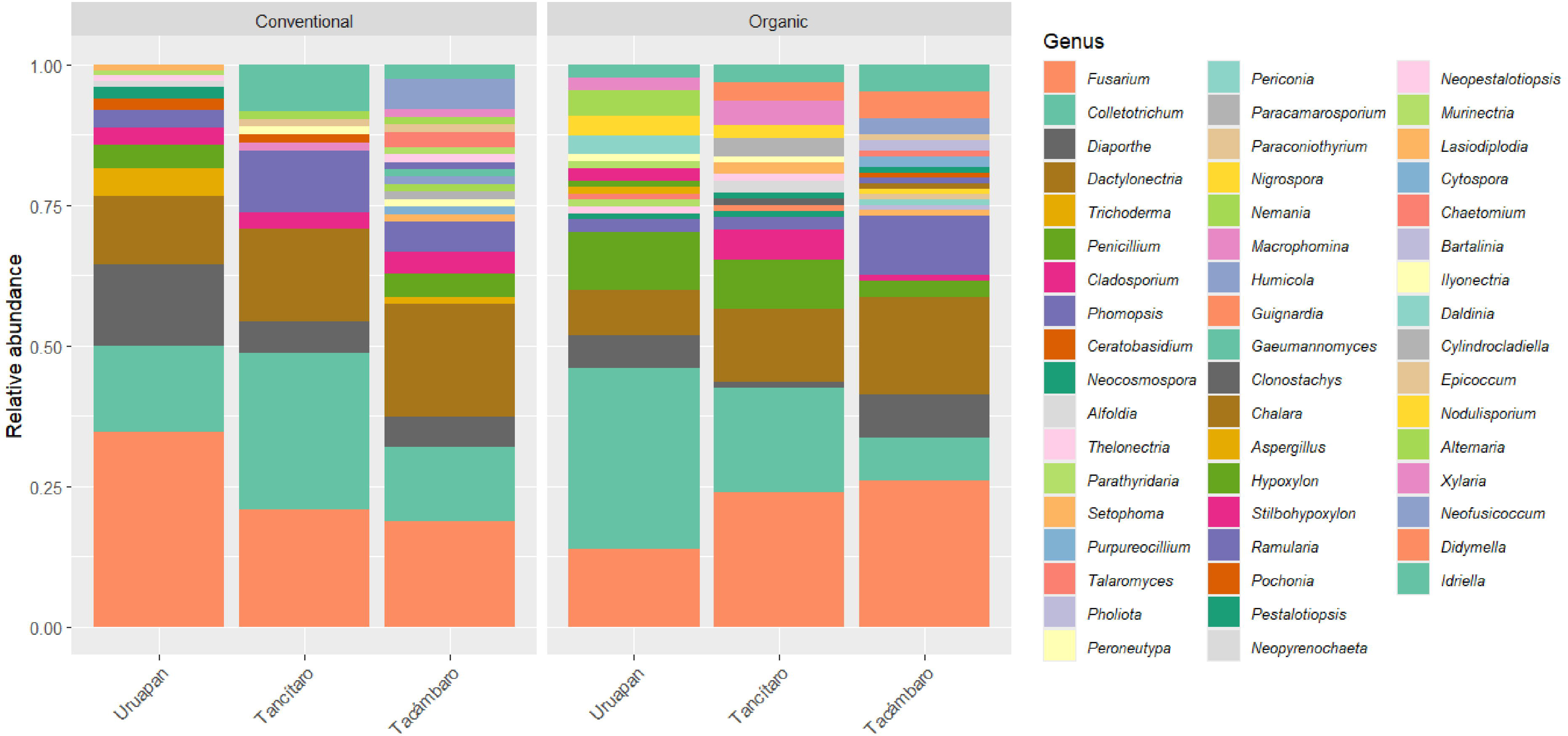
Relative abundance of endophytic fungal genera identified in avocado orchards based on agronomic management and sampling site.

Of the 52 genera identified, 22 were shared between both management systems, while 20 were exclusive to organic orchards and 10 only retrieved in conventional orchards (**Fig. 4a**). Examples of fungal genera exclusively found in organic orchards include *Lasiodiplodia*, *Talaromyces* and *Hypoxylon*, whilst genera such as *Trichoderma* were only found in conventionally managed orchards. In both management systems, Tacámbaro was the county with the largest number of unique fungal genera (12 exclusive genera in conventional orchards and 13 in organic orchards), followed by Uruapan and Tancítaro (**Fig. 4b** and **4c**). Altogether, 17.1 and 16.7 % of fungal genera were common to the three counties in conventional and organic orchards, respectively. Fungal genera found in all orchards were *Colletotrichum*, *Dactylonectria*, *Fusarium* and *Diaporthe* / *Phomopsis.* These genera were also the most abundant, as highlighted in **Fig. 3**.

**Figure 4.**
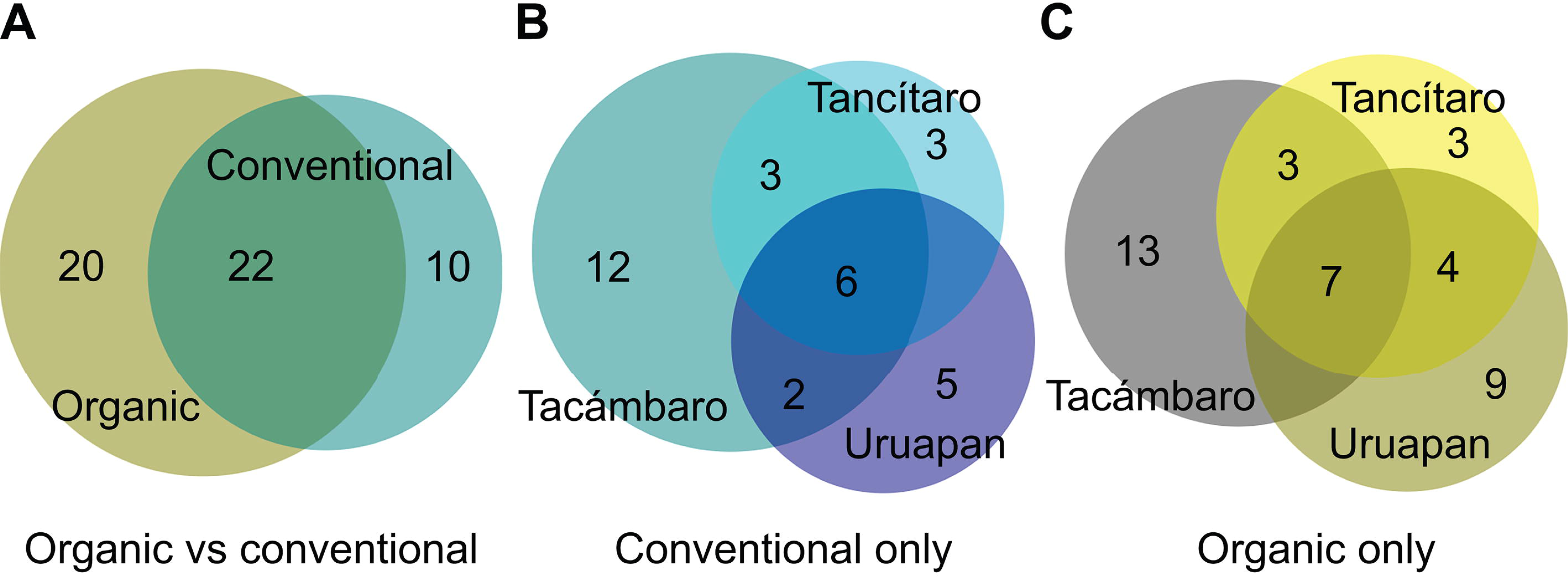
Number of unique and shared endophytic fungal genera. (A) Unique and shared fungal genera between organic and conventional orchards; (B) unique and shared fungal genera within conventional orchards; (C) unique and shared fungal genera within organic orchards.

### Influence of the orchard management type and location on the diversity and community structure of avocado fungal endophytes

Two-way ANOVAs were performed to detect significant differences of management type and sampling site on fungal endophyte species richness (q0), diversity (q1) and evenness (q2), showing that neither site nor the interaction between site and management significantly influenced alpha-diversity metrics (**Supplementary Material S3**). However, the management type significantly affected fungal richness and diversity, both metrics being higher in organic orchards than in conventional orchards (**Fig. 5a and 5b**). On the other hand, the Simpson diversity index (q2) was not significantly influenced by management (**Fig. 5c**), suggesting that, in both systems, dominant species have a similar impact on the structure of culturable endophytic fungal communities associated with avocado trees.

**Figure 5.**
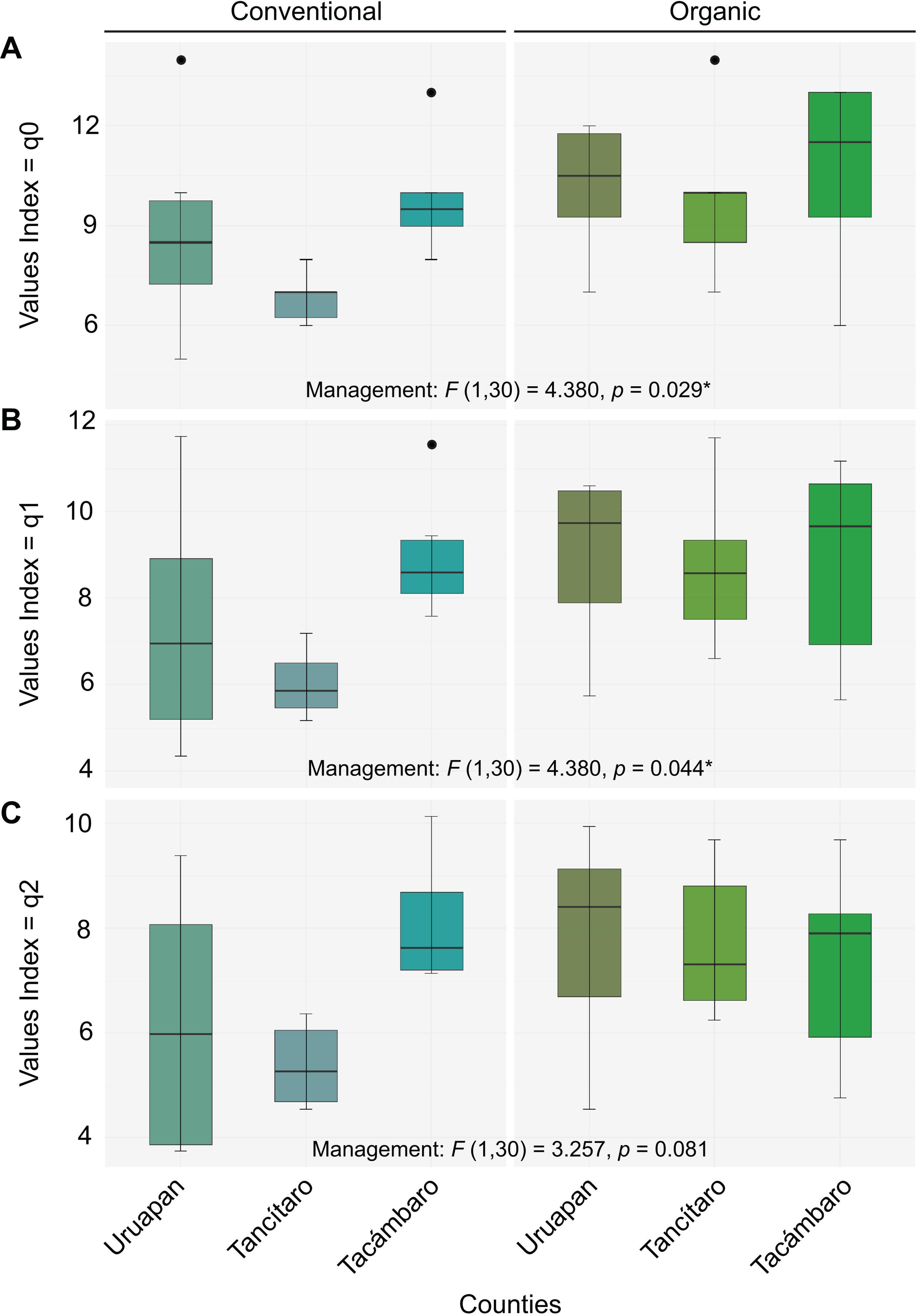
Diversity indices based on Hilĺs numbers (q0: species richness, q1: Shannon diversity, q2: Simpson diversity [dominance]) in conventional and organic orchards from three counties: Uruapan, Tancítaro and Tacámbaro). The boxplots show the median, interquartile range, and outliers for each index and management type. Two-way ANOVA with management type and site as factors showed no significant effect of site nor of the interaction site × management type on alpha-diversity metrics (Supplementary Material S3). Therefore, only results for the effect of management are indicated in the three panels.

Endophytic fungal community structure was analyzed using Non-Metric Multidimensional Scaling (NMDS) based on Bray-Curtis distances. This analysis yielded a *stress value* of *0.14*, indicating an adequate representation of the distances between sites in a two-dimensional space (**Fig. 6**). The results reveal a high degree of similarity in the endophytic fungal community composition associated with avocado trees in Uruapan and Tancítaro, as evidenced by the significant overlap of their ellipses. In contrast, Tacámbaro exhibited a clear separation from these two sites, suggesting a distinctive composition among their samples (**Fig. 6a**). Regarding management practices, a partial separation between conventional and organic systems was observed. Organic communities showed greater dispersion, indicating higher heterogeneity in their composition (**Fig. 6b**). However, when sites (Uruapan, Tancítaro and Tacámbaro) and management practices (conventional and organic) were simultaneously considered, the ellipses showed an overlap within each locality (**Fig. 6c**). The Bray-Curtis dendrogram (**Fig. 6d**) further supports these findings, clearly clustering Uruapan and Tancítaro together, while Tacámbaro appears as a separate group, independently from the management system, highlighting its substantial dissimilarity compared to the other two sites.

**Figure 6.**
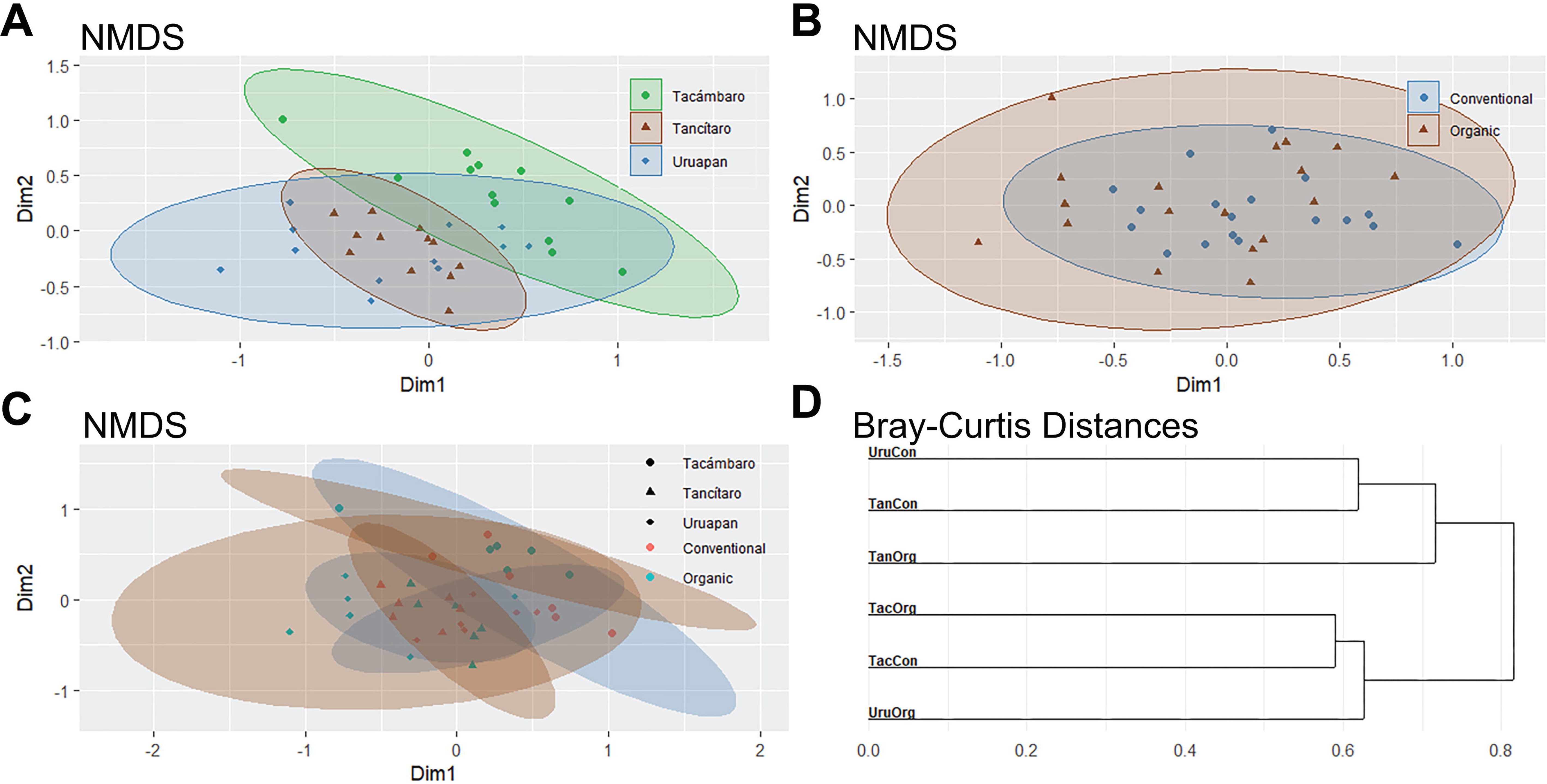
Non-metric multidimensional scaling (NMDS) analysis and Bray-Curtis distance dendrogram for endophytic fungi in avocado orchards. The NMDS plots show sample clustering by (A) site: Uruapan, Tancítaro, and Tacámbaro; (B) agricultural management: conventional and organic; and (C) the combination of both factors. The ellipses represent the 95% confidence interval for each group. (D) The Bray-Curtis dendrogram shows the similarity of fungal communities among the different management systems (Con – Conventional; Org – Organic) and sites (Uru: Uruapan; Tan: Tancítaro; Tac: Tacámbaro).

## Discussion

Despite the economic importance of avocado and the threats that avocado production is facing (soil fertility loss, drought, incidence of pests and diseases), few studies have examined the diversity of avocado fungal endophytes and how they could be impacted by intensive management practices [12]. Considering the benefits that fungal endophytes can provide for their host health and productivity [43, 44], more studies exploring fungal endophytes associated with avocado are crucially needed. Here, we retrieved 529 endophytic fungal isolates from avocado roots and branches, which were grouped into 52 fungal genera, representing 102 tentative species. Most sequenced fungal isolates (98 %) belonged to the Ascomycota phylum, with a dominance of genera such as *Fusarium, Colletotrichum, Dactylonectria,* and *Diaporthe* / *Phomopsis*, which were common to both management types. As these genera comprise species that are known pathogens of avocado [45–48], further research is needed to assess the pathogenicity of these fungal isolates and evaluate whether, under changing environmental conditions, they could switch from endophytes to pathogens [49, 50].

The dominance of *Fusarium* in the fungal endophytic community is not surprising and has been documented in other plants such as vanilla, grapevine and maize [51–53], and recently in avocado nursery plants [12]. *Fusarium, Diaporthe* and *Dactylonectria* have been found to be dominant endophytic genera in chestnut rose [54] or in the weed *Cirsium arvense* [55]. *Phomopsis*, the anamorph state of *Diaporthe*, has been described as an endophyte in rice [56] and several Fabaceae [57, 58], among many other hosts. *Colletotrichum* species have also been described as endophytes in apple and tomato, among other crops [59, 60]. The predominance of *Colletotrichum* in the fungal endophytic community was recently reported in the invasive plant *Ageratina adenophora* in China [61], thus raising concerns about the possible spill-over to significant crops in the region. In Michoacán, the avocado producing region where our sampling took place, a possible spill-over of latent pathogens to neighboring plant species could threaten the integrity of pine-oak forests surrounding orchards, which are already at risk due to the pressure exerted by avocado expansion [62]. For example, *Fusarium circinatum*, a species tentatively identified in our study (although its taxonomic assignment should be refined with other molecular markers), is an important pine pathogen [63]. *Neofusicoccum,* a genus exclusively detected in Tacámbaro county, has been determined as the causal agent of pine ghost canker in Southern California [64]. Some *Colletotrichum* species have been associated with oak anthracnose [65], *Phomopsis / Diaporthe* with oak cankers [66], and species of *Colletotrichum, Diaporthe* and *Ilyonectria* with oak seedling dieback, thereby limiting forest natural regeneration [67]. Mexico is a center of diversity both for pine and oak species [68] and considering the key role these forests will play in mitigating climate change, preserving their health and productivity is paramount. Concrete efforts should thus be made in assessing the risk that these latent pathogens may represent for increasingly fragmented forests surrounding avocado orchards.

Our study revealed that endophytic fungal richness and diversity were higher in organic avocado orchards than in those conventionally managed. The positive influence of organic management on the diversity of fungal endophytes has been previously shown in annual crops, such as soybean, pepper, melon and maize [19, 20]. Although less investigated, organic management has also been reported to enhance endophytic fungal richness in perennial crops such as grapevine [69]. Our study thus confirms the positive effect of organic management on the richness and diversity of avocado-associated endophytic fungi, adding further evidence to the large body of literature highlighting the negative impact of intensive agricultural management on crop-associated endophytic fungi.

In contrast with a previous report [28], no significant effect of management practices was detected on the endophytic fungal community structure. However, Venn diagrams showed that only 42% of fungal taxa were shared between conventional and organic orchards, which indicates an effect of management practices on the endophytic fungal community composition. For example, the genus *Trichoderma* was only detected in conventional orchards. The benefits of *Trichoderma* have been widely highlighted, as a plant growth promoter, plant immunity primer, and antifungal and entomopathogenic agent [70]. Conversely, taxa such as *Hypoxylon* and *Talaromyces* were only retrieved in organic orchards. The genus *Hypoxylon* has been described for its potential in biocontrol strategies, due to the strong bioactivity of its metabolites [71, 72]. Plant beneficial effects of endophytic *Talaromyces* were also reported, for plant nutrient acquisition and protection against pathogens [73, 74]. As mentioned earlier, most reports on avocado endophytes have been focused on the genus *Trichoderma*, which only represented 1.1 % of the retrieved isolates in our study. Our findings thus highlight the need to harness the biotechnological potential of less studied endophytic fungal taxa associated with this tree crop, which could also contribute to orchard sustainable management practices.

Finally, our results revealed a stronger impact of site than of management practices on beta-diversity metrics, suggesting the role of local factors in shaping fungal endophytic communities. This is supported by the Venn diagrams showing that, within a management type, few taxa were shared between sites. The importance of the local environment on the structure of root endophytic fungal communities was demonstrated at the continental scale for the Brassicaceae *Microthlaspi* [75], evidencing the role of climatic variables as drivers of the endophytic community composition. The significant influence of geographical distance on fungal endophytes was also shown in poplar, agave, and oak [76–78] and was attributed to the dispersal limitation of fungal spores [79]. In our work, the endophytic fungal community of Tacámbaro orchards were separated from those of Uruapan and Tancítaro, independently from the orchard management type. Although climate in the three counties is classified as sub-humid temperate, Tacámbaro shows a narrower range of mean annual precipitation (900 to 1300 mm) than the other two counties (700 – 2000 mm) [80]. Other environmental factors should be examined in detail to determine the potential drivers of the endophytic fungal community composition, especially because Tacámbaro was recently identified as a future hotspot of avocado expansion [81]. These findings confirm the importance of local factors as determinants of endophytic fungal assemblages.

### Conclusions

Our results show that culturable endophytic fungi associated with avocado branches and roots respond to both geographic site and agricultural management. Conventional management practices decreased the richness and diversity of avocado endophytic fungi, although fungal community structure was mainly driven by factors inherent to orchard location. Dominant endophytic genera were common to all orchards, regardless of management type or site, and represent potential avocado pathogens. Other possibly detected fungal species comprise known pathogens of pines and oaks and may represent a threat for the forests at the avocado expansion limit. A critical assessment of the environmental conditions under which fungal endophytes may switch to avocado or forest tree pathogens is therefore required. Conversely, under the current scenario of avocado cultivation in the state of Michoacán and its multiple adverse consequences, the biotechnological potential of beneficial endophytic fungi needs to be fully assessed, as these isolates may be promising candidates for the development of novel bioformulations.

## Acknowledgements

F.R. was supported by a CONAHCYT-PRONACES-PPE grant (project “PERSEA”, project number 322772). D.S.H. is grateful for his CONAHCYT postgraduate fellowship. The authors express their gratitude to the avocado producers who generously granted access to their orchards; Enrique Bautista Villegas, Gabriel Guzman Pardo, and Abraham Pardo Cuevas. We also extend our thanks to Edgar Guevara-Avendaño for his assistance with DNA extraction protocols, Edith Garay-Serrano for her help with fungi molecular identification, Rosaura Guadalupe Alfaro-García for her help with the calculation of Hill numbers, and Alfonso Méndez-Bravo for his assistance with artwork editing.

## Declaration of competing interests

The authors declare no competing interests.

## Data availability statement

The datasets generated during and/or analysed during the current study are available from the corresponding author on reasonable request.

## Notes

### Competing Interest Statement

The authors have declared no competing interest.

