## Supplementary Material S1 for "Diversity of culturable endophytic fungi associated with avocado orchards under organic and conventional management"

**Supplementary Material S1.** Localization of sampled orchards. Left panel: geographic projection of Mexico with the state of Michoacán highlighted in dark green. Right panel: localization of sampled orchards in the counties of Uruapan (El Capulin, El Paraíso), Tancítaro (El Cerrito de Varal, Palo Amarillo) and Tacámbaro (María, La Tiendita 1). The table describes the management type of each sampled orchard, by county.

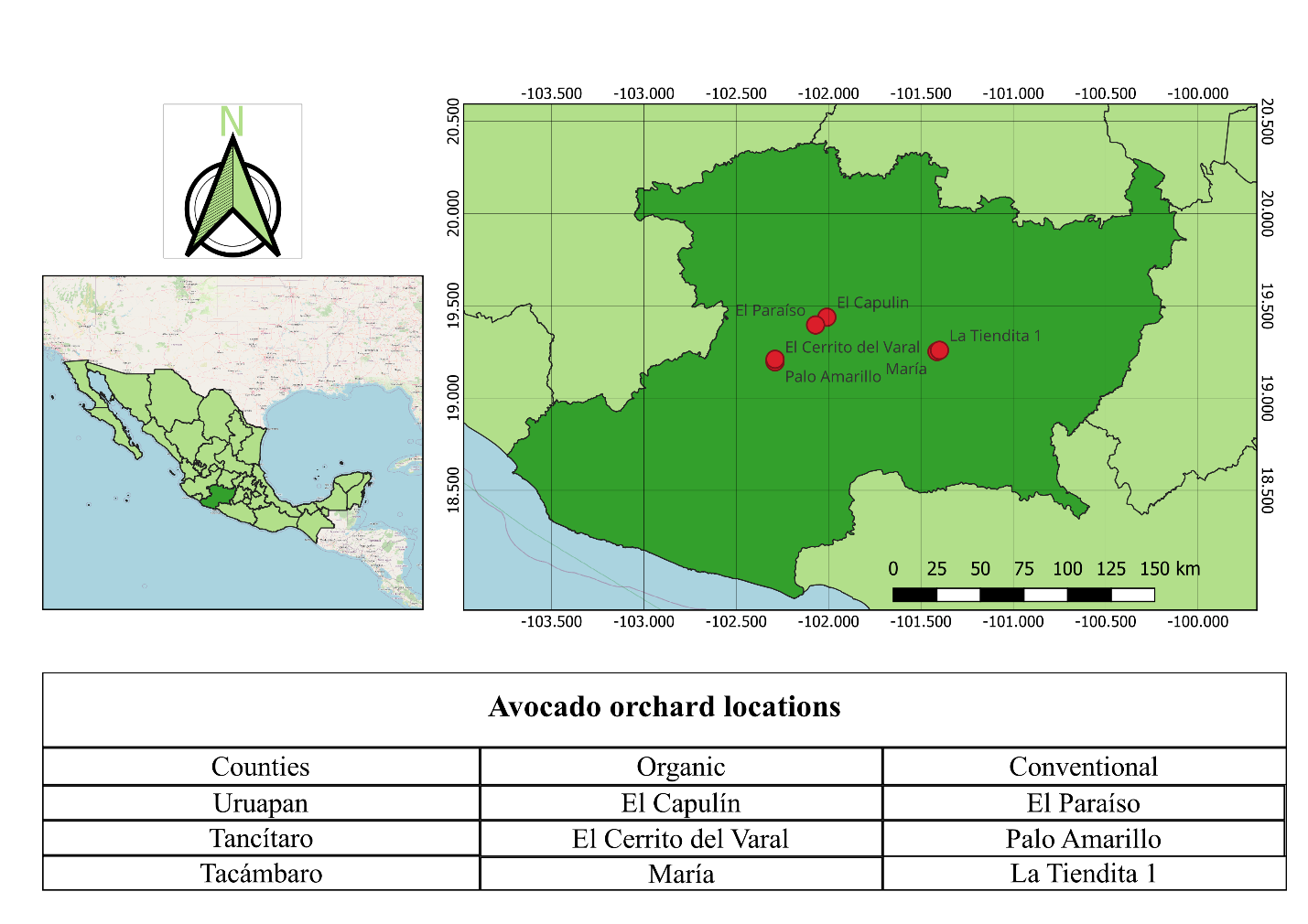
