## Supplementary Material S3 for "Diversity of culturable endophytic fungi associated with avocado orchards under organic and conventional management"

**Supplementary Material S3.** Two-way ANOVA results for alpha-diversity metric of the endophytic fungal community with management type and sampling site as factors

|  |  | **Df** | **Sum Sq** | **Mean Sq** | **F value** | ***p* value** |
| --- | --- | --- | --- | --- | --- | --- |
| **Richness (q0)** | Management | 1 | 26.69 | 26.694 | 5.127 | **0.0296 *** |
|  | Site | 2 | 22.39 | 11.194 | 2.188 | 0.1297 |
|  | Management × Site | 2 | 7.72 | 3.861 | 0.755 | 0.4789 |
|  | Residuals | 30 | 153.50 | 5.117 |  |  |
| **Diversity (q1)** | Management | 1 | 17.36 | 17.361 | 4.384 | **0.0488 *** |
|  | Site | 2 | 14.51 | 7.257 | 1.833 | 0.1775 |
|  | Management × Site | 2 | 12.06 | 6.029 | 1.522 | 0.2346 |
|  | Residuals | 30 | 118.80 | 3.690 |  |  |
| **Evenness (q2)** | Management | 1 | 10.09 | 10.091 | 3.257 | 0.0812 |
|  | Site | 2 | 8.31 | 4.157 | 1.342 | 0.2766 |
|  | Management × Site | 2 | 16.00 | 8.000 | 2.582 | 0.0923 |
|  | Residuals | 30 | 92.95 | 3.098 |  |  |
